# UniqTag: Content-derived unique and stable identifiers for gene annotation

**DOI:** 10.1101/007583

**Authors:** Shaun Jackman, Joerg Bohlmann, Inanҫ Birol

## Abstract

**Summary:** When working on an ongoing genome sequencing and assembly project, it is rather inconvenient when gene identifiers change from one build of the assembly to the next. The gene labelling system described here, UniqTag, addresses this common challenge. UniqTag assigns a unique identifier to each gene that is a representative *k* - mer, a string of length *k*, selected from the sequence of that gene. Unlike serial numbers, these identifiers are stable between different assemblies and annotations of the same data without requiring that previous annotations be lifted over by sequence alignment. We assign UniqTag identifiers to nine builds of the Ensembl human genome spanning seven years to demonstrate this stability.

**Availability and implementation:** The implementation of UniqTag is available at https://github.com/sjackman/uniqtag

Supplementary data and code to reproduce it is available at https://github.com/sjackman/uniqtag-paper

## 1 INTRODUCTION

The task of annotating the genes of a genome sequence often follows genome assembly. These annotated genes are assigned unique identifiers by which they can be referenced. Assembly and annotation is frequently an iterative process, by refining the method or by the addition of more sequencing data. These gene identifiers would ideally be stable from one assembly and annotation to the next. The common practice is to use serial numbers to identify genes that are annotated by software such as MAKER (Campbell, 2014), which, although certainly unique, are not stable between assemblies. A single change in the assembly can result in a total renumbering of the annotated genes.

One solution to stabilize identifiers is to assign them based on the content of the gene sequence. A cryptographic hash function such as SHA (Secure Hash Algorithm) (Dang, 2012) derives a message digest from the sequence, such that two sequences with the same content will have the same message digest, and two sequences that differ will have different message digests. If a cryptographic hash were used to identify a gene, the same gene in two assemblies with identical content would be assigned identical identifiers. Yet, by design a slight change in the sequence, such as a single-character substitution, would result in a completely different digest.

Locality-sensitive hashing in contrast aims to assign items that are similar to the same hash value. A hash function that assigns an identical identifier to a sequence after a modification of that sequence is desirable for labelling the genes of an ongoing genome annotation project. One such locality-sensitive hash function, MinHash, was employed to identify web pages with similar content (Broder, 1997) by selecting a small representative set of words from a web page.

UniqTag is inspired by MinHash. It selects a single representative *k*-mer from a sequence to assign a stable identifier to a gene. These identifiers are intended for systematic labelling of genes rather than assigning biological gene names, as the latter are typically based on biological function or homology to orthologous genes.

## 2 METHODS

The UniqTag is defined mathematically as follows. Let Σ be an alphabet, such as the twenty standard amino acids or the four nucleotides. Let Σ*^k^* be the set of all strings over Σ of length *k*. Let *s* be a string over Σ, such as the peptide or nucleotide sequence of a gene. Let *C*(*s*) be the set of all substrings of *s*, and *C_k_* (*s*) be the set of all *k*-mers of *s*, that is, all substrings of *s* with length *k*.

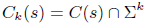

Let *S* be a set of *n* strings {*s*_0_, …, *s*_n_}, such as the peptide or nucleotide sequences of the annotated genes of a genome assembly. Let *f*(*t, S*) be the frequency in *S* of a *k*-mer *t*, defined as the number of strings in *S* that contain the *k*-mer *t*.

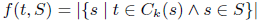

Let *T* be the set of *k*-mers of *t*, and min *T* be the lexicographically minimal *k*-mer of *T*. If the *k*-mers of *T* were sorted alphabetically, it would be the first *k*-mer in the list.

Finally, *u_k_* (*s, S*) is the UniqTag, the lexicographically minimal *k*-mer of those *k*-mers of *s* that are least frequent in *S*.

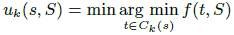

Typically, *u_k_* (*s*, *S*) is the first *k*-mer in an alphabetically sorted list of the *k*-mers of a gene that are unique to that gene.

A UniqTag can be generated from the nucleotide sequence of a gene or the translated peptide sequence of a protein-coding gene. Using the peptide sequence results in a UniqTag that is stable across synonymous changes to the coding sequence as well as to changes in the untranslated regions and introns of the gene. Since the amino acid alphabet is larger than the nucleotide alphabet, fewer characters are required for a *k*-mer to be likely unique, resulting in an aesthetically pleasing shorter identifier.

When two gene models have identical *k*-mer compositions, they would be assigned the same UniqTag. It is also possible that two genes that have no unique *k*-mer and similar *k*-mer composition are assigned the same UniqTag. In such cases, genes that have the same UniqTag are distinguished by adding a numerical suffix to the UniqTag.

The UniqTag is designed to be stable but will change in the following conditions: (1) when the sequence at the locus of the UniqTag changes; (2) when a least-frequent *k*-mer that is lexicographically smaller than the previous UniqTag is created; (3) when a duplicate *k*-mer is created elsewhere that results in the previous UniqTag no longer being a least-frequent *k*-mer.

The special cases of merging and splitting gene models are interesting. Concatenating two gene models results in a gene whose UniqTag is the minimum of the two previous UniqTags, unless the new UniqTag spans the junction of the two sequences. Similarly when a gene model is split in two, one gene is assigned a new UniqTag and the other retains the previous UniqTag, unless the previous UniqTag spanned the junction.

Importantly and in contrast, unlike naming the genes after the genomic contigs or scaffolds in which they are found, changing the order of the genes in a genome assembly has no effect on the UniqTag.

## 3 RESULTS

To demonstrate the stability and utility of UniqTag, we assigned identifiers to the genes of nine builds of the Ensembl human genome (Flicek, 2014) spanning seven years and two major genome assemblies, NCBI36 up to build 54 and GRCh37 afterward. An identifier of nine peptides (*k* = 9) was assigned to the first protein sequence, that with the smallest Ensembl protein (ENSP) accession number, of each gene. The number of common UniqTag identifiers between older builds from build 40 on and the current build 75 is shown in Figure 1. Also shown is the number of common gene and protein identifiers (ENSG and ENSP accession numbers) between builds and the number of genes with peptide sequences that are identical between builds. Although less stable than the gene ID, the UniqTag is more stable than the protein ID and the peptide sequence.

**Fig. 1.**
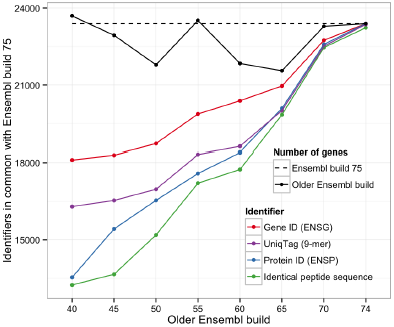
The number of common UniqTag identifiers between older builds of the Ensembl human genome and the current build 75, the number of common gene and protein identifiers between builds, and the number of genes with peptide sequences that are identical between builds.

Whereas the gene and protein identifiers can, with effort, be lifted over from older builds to the newest build, the UniqTag identifier can be generated without any knowledge of previous assemblies, making it a much simpler operation. The number of identical peptide sequences between builds shows the stability that would be expected of using a cryptographic hash value of the peptide sequence as the identifier. Supplementary Figure S1 shows that the UniqTag stability is insensitive to the size of the UniqTag identifier for values of *k* between 8 and 50 peptides. The data for these figures are shown in supplementary Table S1.

## ACKNOWLEDGEMENTS

The authors thank Nathaniel Street for his enthusiastic feedback, the SMarTForests project and the organizers of the 2014 Conifer Genome Summit that made our conversation possible.

